# A Humanized Mouse Model of Oral Squamous Cell Carcinoma for Investigating Resistance to Anti-PD1 Immunotherapy

**DOI:** 10.1101/2025.06.13.659630

**Authors:** Amit Kumar Chakraborty, Chumki Choudhury, Rajnikant Dilip Raut, Manish V Bais

**Affiliations:** Department of Translational Medicine, Boston University School of Medicine, Boston, MA

**Keywords:** Humanized, OSCC, Anti-PD1, Pembrolizumab, Resistant, HNSCC

## Abstract

Immunotherapy has transformed cancer therapeutics; however, its impact on oral squamous cell carcinoma (OSCC) remains limited due to acquired resistance, including to Pembrolizumab. To better understand this challenge, we develop a humanized mouse model using immune-deficient NCG mice engrafted with human peripheral blood mononuclear cells (hPBMC), followed by orthotopic implantation of head and neck squamous cell carcinoma (HNSCC) stem cells. This model closely mimics the human tumor-immune microenvironment, showing aggressive tumor growth and metastasis. Pembrolizumab treatment significantly decreases CD8+ T cells, dendritic cells, and MHC-class-I expression at tumor sites, which are related to immunotherapy resistance. Further, pembrolizumab treatment confirms the elevated PD-L1 levels, immune evasion pathways and downregulation of MHC-class-I. qPCR further validates human-specific HLA expression. Our model offers a valuable tool for studying immunotherapy resistance and advancing treatment strategies in OSCC and could be useful for other cancers.

## Introduction

Oral squamous cell carcinoma (OSCC), a subset of head and neck squamous cell carcinoma (HNSCC), remains a major global health challenge owing to its aggressive clinical course, high recurrence rates, and limited responsiveness to conventional therapies.^1,2^ The advent of immune checkpoint inhibitors (ICIs), particularly anti-programmed cell death protein 1 (PD-1) antibodies, such as Pembrolizumab and Nivolumab, has significantly advanced treatment paradigms by enhancing host antitumor immunity.^3,4^ However, a substantial proportion of patients exhibit primary or acquired resistance to ICIs, highlighting the urgent need to elucidate the underlying mechanisms of therapeutic failure and to identify strategies to overcome resistance.^5^

Patients partially respond to current immunotherapies, with immune-excluded and immune-desert (“cold”) tumors representing the majority of non-responders.^6^ The immune landscape of tumors, including permissive or restrictive, plays a critical role in determining T cell activity and therapeutic success or failure.^7–11^ Thus, future therapies need detailed mechanistic insight. A deeper understanding of both tumor-intrinsic and-extrinsic mechanisms of immune escape, such as MHC-I loss, neoantigen depletion, and suppressive cell populations, is essential. However, lack of specific models hampers progress. Developing an immunotherapy-resistant model, especially a humanized cancer model, could provide support to the next generation of immunotherapy to benefit therapeutic-resistant patient tumors.

Humanized mouse models, generated through engraftment of human peripheral blood mononuclear cells (PBMCs) into immunodeficient hosts, offer a powerful platform for studying human immune responses *in vivo*.^12^ This model recapitulates key features of the human tumor-immune microenvironment, enabling the mechanistic investigation of immune evasion, checkpoint blockade resistance, and immune-related adverse events (irAEs). Notably, irAEs, which are paradoxical immune responses triggered by ICIs, can exacerbate disease progression and are increasingly being recognized as clinically relevant phenomena in patients receiving anti-PD-1 therapy.^13,14^

In the present study, we established and rigorously validated a humanized mouse model of OSCC by orthotopically implanting patient-derived HNSCC stem cells into PBMC-engrafted NCG mice. This dual engraftment approach enabled the formation of a clinically significant tumor immune microenvironment. We comprehensively characterized immune cell infiltration, tumor progression, and transcriptomic alterations in response to pembrolizumab treatment. Our findings revealed paradoxical tumor-promoting effects of anti-PD-1 therapy, accompanied by depletion of CD8^+^ T cells and dendritic cells, upregulation of PD-L1, and regulation of immune resistance mechanisms, including MHC class I dysregulation.^15^

Importantly, we validated the human-specific expression of key immune markers, including HLA-A and HLA-B (MHC class I molecules) to confirm the immunological fidelity of the model. By capturing both the therapeutic responses and resistance mechanisms, including those resembling irAEs, this model provides a robust and translationally relevant platform for dissecting immune dynamics and guiding the development of next-generation immunotherapies for OSCC and related malignancies.

## Results

### Establishment and Validation of a Humanized Mouse Model

To develop a model for studying human immune responses in OSCC, we engrafted immunocompromised NCG mice with hPBMCs via tail vein injection (Fig. 1a). We assessed the distribution and infiltration of human immune cells in the tongue and spleen. Flow cytometric analysis detected human CD45^+^ immune cells (Fig. 1b). As expected, CD45^+^, CD4^+^, CD8^+^ T cells, and conventional DC populations were increased in mice tongues injected with hPBMC, thereby capturing the heterogeneity of human immune infiltration in the tongue (Fig. 1c, d). Notably, PD-1^+^ CD8^+^ T cells were also enriched in the engrafted mice, indicating an active immune microenvironment (Fig. 1d). Further validation of immune reconstitution showed a significant increase in human CD4^+^, CD3^+^, CD4^+^, and CD8^+^ T cells in the spleen (Fig. 1e, f). High-dimensional immune profiling using opt-SNE projections for flow cytometry data revealed distinct clustering of major immune subsets, including CD45^+^, CD4^+^, and CD8^+^ T cells, and conventional DC populations, thereby capturing the heterogeneity of the human immune infiltrate (Fig. 1c, e). Immunofluorescence staining of tongue sections revealed distinct localization of HLA-A+ human immune cells in hPBMC-engrafted mice, with minimal signal in non-engrafted controls (Fig. 1g), confirming successful tissue-level humanization. Together, these data confirm the successful establishment of a functional human immune compartment in NCG mice.

**Figure 1:**
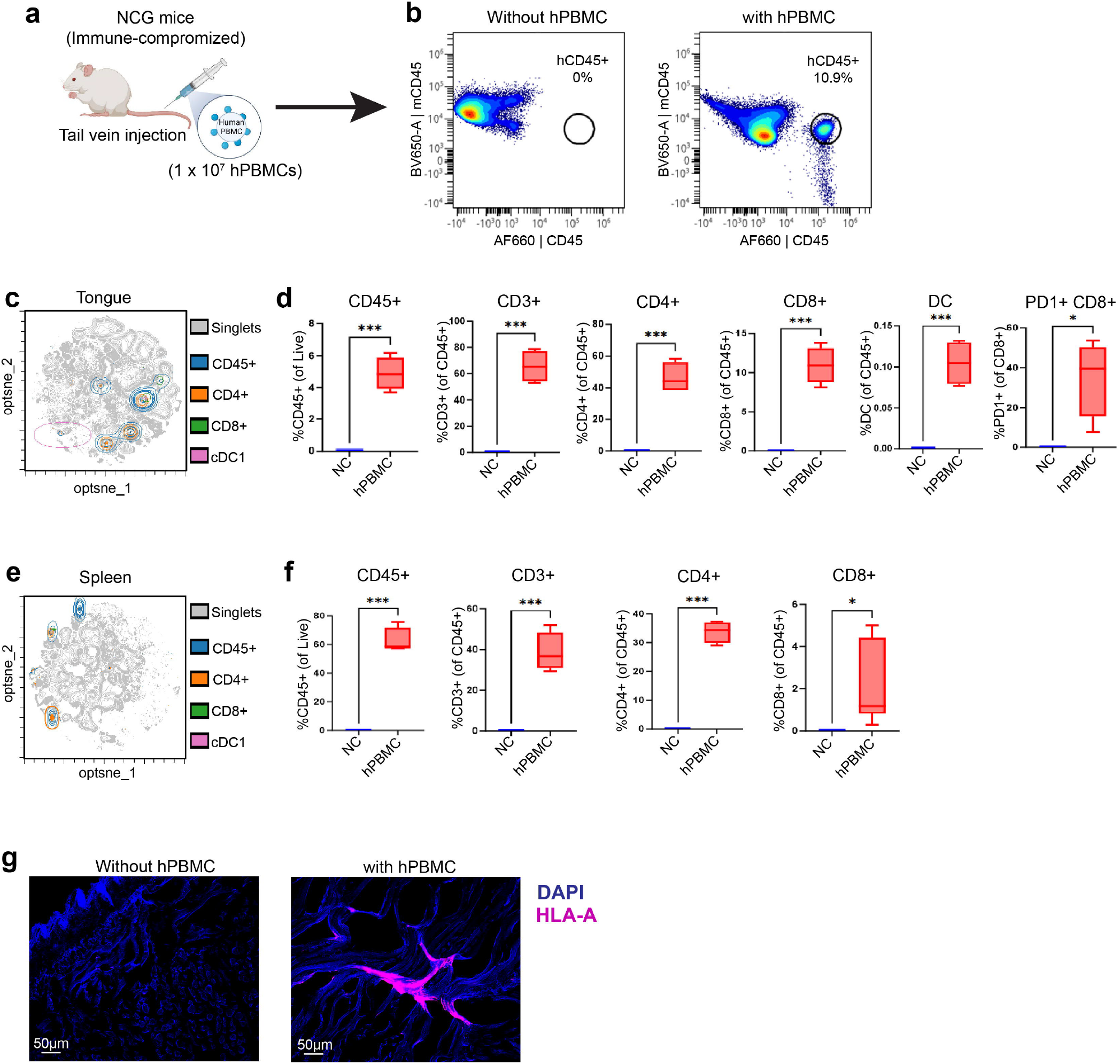
Establishment and validation of a humanized mouse model. **a)** Schematic overview of hPBMC engraftment into NCG mice. **b)** Flow cytometry gating strategy confirming successful engraftment, with detection of human CD45+ leukocytes in peripheral blood. **c)** opt-SNE projection of flow cytometry data showing distinct clusters of human T cells and dendritic cells (DCs) in the mouse tongue. **d)** Expression of major human immune cell markers (e.g., CD45, CD3, CD4, CD8, CD11c) were detected in the tongue, confirming local immune cell infiltration. **e)** opt-SNE analysis of spleen samples reveals robust presence of human T cells and DCs. **f)** Quantification of human immune cell populations in the spleen by flow cytometry, showing significant expansion post-engraftment. **g)** Immunofluorescence staining for human HLA-A in mouse tissues demonstrates spatial localization and integration of human immune cells, supporting successful humanization

### Orthotopic Implantation of HNSCC Stem Cells in Humanized Mice

To evaluate tumor-immune interactions in humanized mice, we orthotopically implanted human HNSCC stem cells into the tongues of hPBMC-engrafted NCG mice, resulting in tumor formation and enabling the study of human immune responses within a tumor-bearing microenvironment (Fig. 2a). Flow cytometry analysis revealed substantial infiltration of human immune cells into both tongue (Fig. 2b) and spleen tissues (Fig. 2c). Specifically, we observed significant increases in CD45^+^, CD3+, CD4 +, CD8 + T cells, and dendritic cells compared with hPBMC only, indicating active immune surveillance in the tongue tumor (Fig. 2d, e). Interestingly, the levels of CD45+ immune cells in the spleen were significantly reduced, along with a noticeable reduction in CD3 +, CD4 +, and CD8 + T cells, suggesting their active localization to the tumor site (Fig. 2f, g). High-dimensional opt-SNE projections further illustrated the complexity of the immune landscape, with clear clustering of CD45 +, CD4 +, and CD8 + T cell populations in both the tongue and spleen tissues (Fig. 2d, f). Immunofluorescence imaging using DAPI and anti-HLA-A antibodies confirmed the presence and localization of hPBMCs within the tumor site, validating the establishment of a functional human immune compartment in tumor-bearing mice (Fig. 2h). Together, these findings demonstrate that the orthotopic implantation of HNSCC stem cells into humanized mice generates a dynamic and responsive tumor-immune microenvironment. This model provides a valuable platform for studying human immune responses to solid tumors and for the preclinical evaluation of immunotherapeutic strategies.

**Figure 2:**
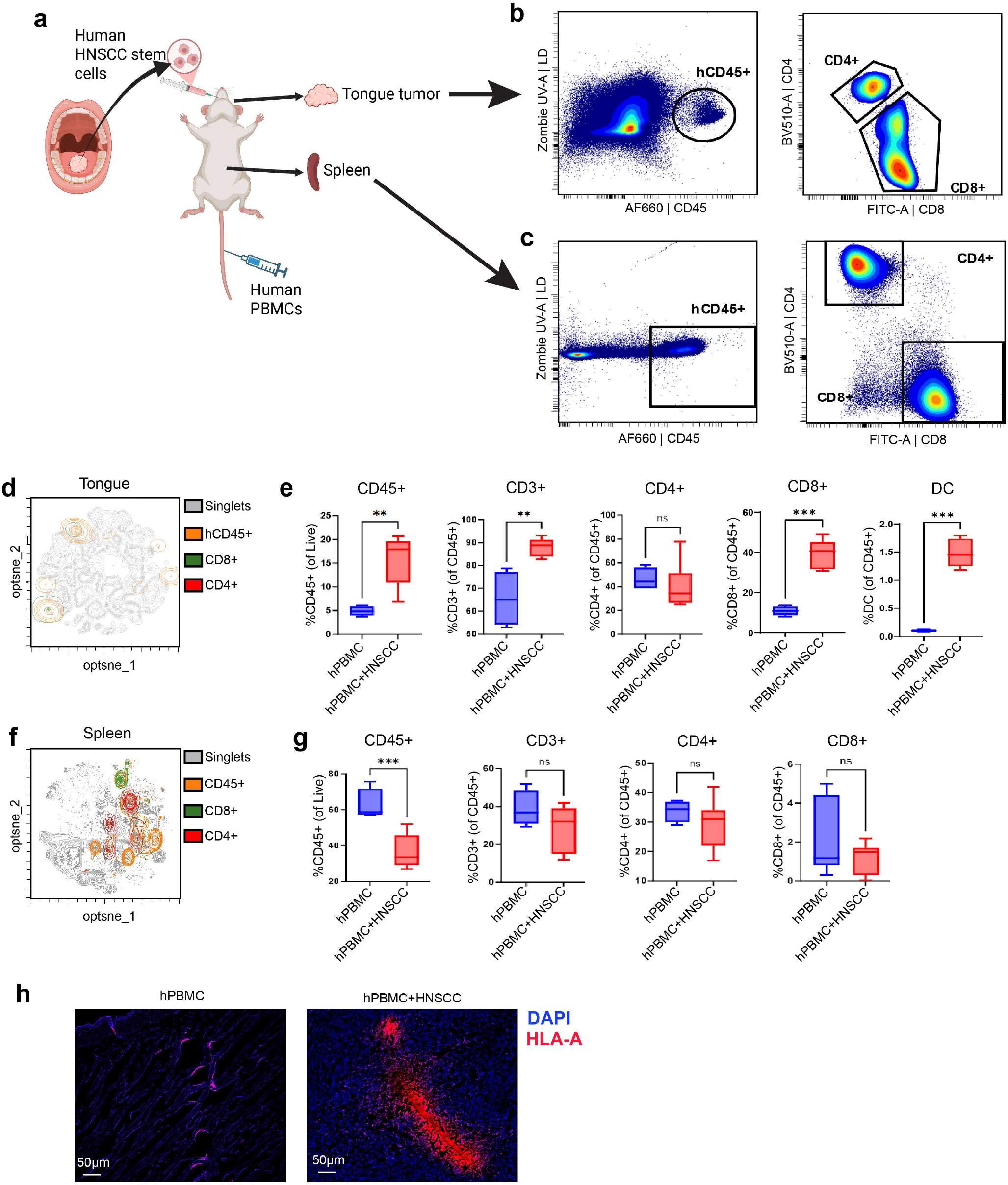
Orthotopic implantation of human HNSCC stem cells in humanized mice. **a)** Schematic representation of the orthotopic injection of human head and neck squamous cell carcinoma (HNSCC) stem cells into the tongues of NCG mice previously engrafted with human PBMCs, establishing a humanized tumor model. **b)** Flow cytometry gating strategy confirming the presence of human CD45+ immune cells in the tumor-bearing mouse tongue. **c)** Detection of human CD45+ immune cells in the spleen, indicating systemic immune cell distribution. **d)** opt-SNE projection of flow cytometry data showing distinct populations of human T cells and dendritic cells (DCs) infiltrating the tongue tumor microenvironment. **e)** Quantification of immune cell infiltration reveals a significant increase in human immune cells within tongue tumors compared to controls (p < 0.01). **f)** opt-SNE analysis of spleen samples showing stable populations of human T cells and DCs, consistent with systemic immune reconstitution. **g)** Comparative analysis indicates that the immune landscape in the spleen remains largely unchanged following tumor implantation, suggesting localized immune modulation within the tumor site.

### Tumor metastasis following anti-PD-1 therapy

To evaluate the therapeutic effect of anti-PD-1 immunotherapy with pembrolizumab (αPD-1), we treated hPBMC-engrafted mice bearing orthotopic HNSCC tumors with pembrolizumab or immunoglobulin G (IgG) as controls (Fig. 3a). Mice receiving both αPD-1 therapy and IgG exhibited a significant tumor burden, with no significant difference between the groups (Fig. 3b). Histological analysis of the tongue revealed similar numbers of tumor lesions in both the IgG and αPD-1 treated groups in comparison with hPBMC only mice (Fig. 3c). Tumor lesions were also detected in the livers of IgG-as αPD-1 treated mice (Fig. 3d and e). Surprisingly, liver metastasis was significantly higher in αPD-1 treated mice compared to IgG-treated mice (Fig. 3e). Histological analysis further validated the presence of tumor lesions in higher numbers in αPD-1 treated mice compared to IgG-treated mice (Fig. 3f, h). HLA-A staining in the liver showed a significant increase in hPBMCs (HLA-A) in the IgG-treated mice group compared to that in the hPBMC-only mice group. Surprisingly, HLA-A intensity decreased in αPD-1 treated mice compared to that in the IgG-treated group, suggesting less infiltration of hPBMCs into the tumor site (Fig. 3g). Together, these data suggest that tumor metastasis to the liver was accelerated after αPD-1 treatment, with decreased immune infiltration at the tumor site.

**Figure 3:**
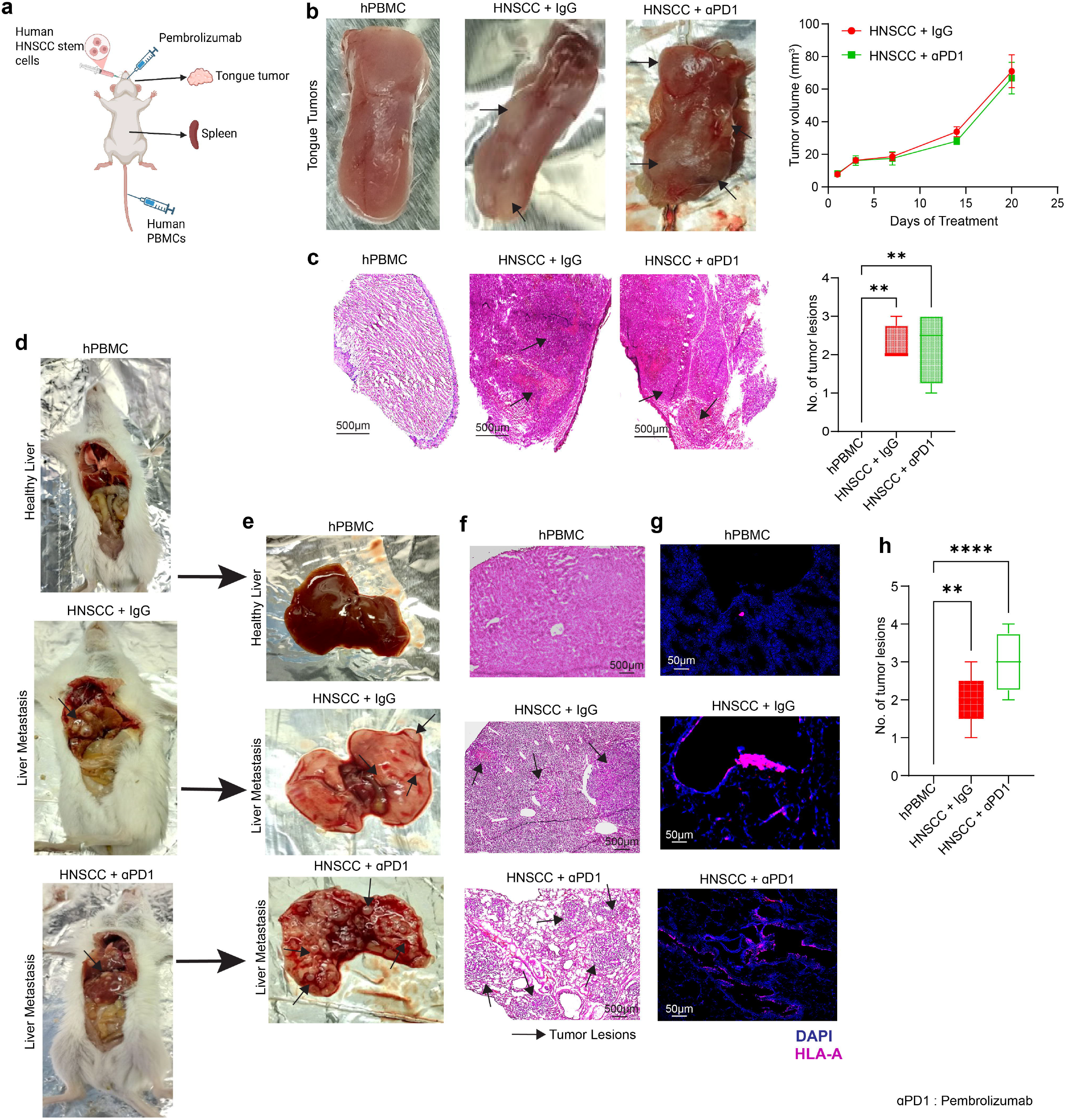
Tumor progression and metastatic spread following Pembrolizumab treatment in humanized mice. **a)** Schematic representation of Pembrolizumab administration in humanized mice bearing orthotopic tongue tumors derived from human HNSCC stem cells. **b)** Gross images of tongue tumors with quantification of tumor volume (mm^3^) showing increased tumor growth following Pembrolizumab treatment. **c)** Hematoxylin and eosin (H&E) staining of tongue sections illustrating tumor lesions, with comparative analysis of lesion size post-treatment. **d-e)** Representative images of gross liver metastases in humanized mice following HNSCC stem cell implantation and Pembrolizumab treatment. **f)** H&E staining of liver sections confirming metastatic tumor lesions. **g)** Immunofluorescence staining for human HLA-A in liver tissue, demonstrating human immune cell presence. **h)** Quantification of liver tumor lesions reveals an increased metastatic burden in Pembrolizumab-treated mice.

### Validation of αPD-1 treatment Therapy Resistance

To determine the immune profile following αPD-1 therapy, we performed a flow cytometry analysis (Fig. 4a). This analysis showed a notable decrease in important immune cell populations within the tongue tumor microenvironment, such as CD45+, CD3+, CD8^+^ T cells, and DCs (Fig. 4b), in contrast to the IgG-treated group. In the spleen, we noted a significant reduction in both CD4^+^ and CD8^+^ T cells after αPD-1 treatment, whereas the total CD45 + immune cell count remained unchanged (Fig. 4c). This reduction in immune cells suggests overall suppression of the immune response and resistance to αPD-1 treatment.

**Figure 4:**
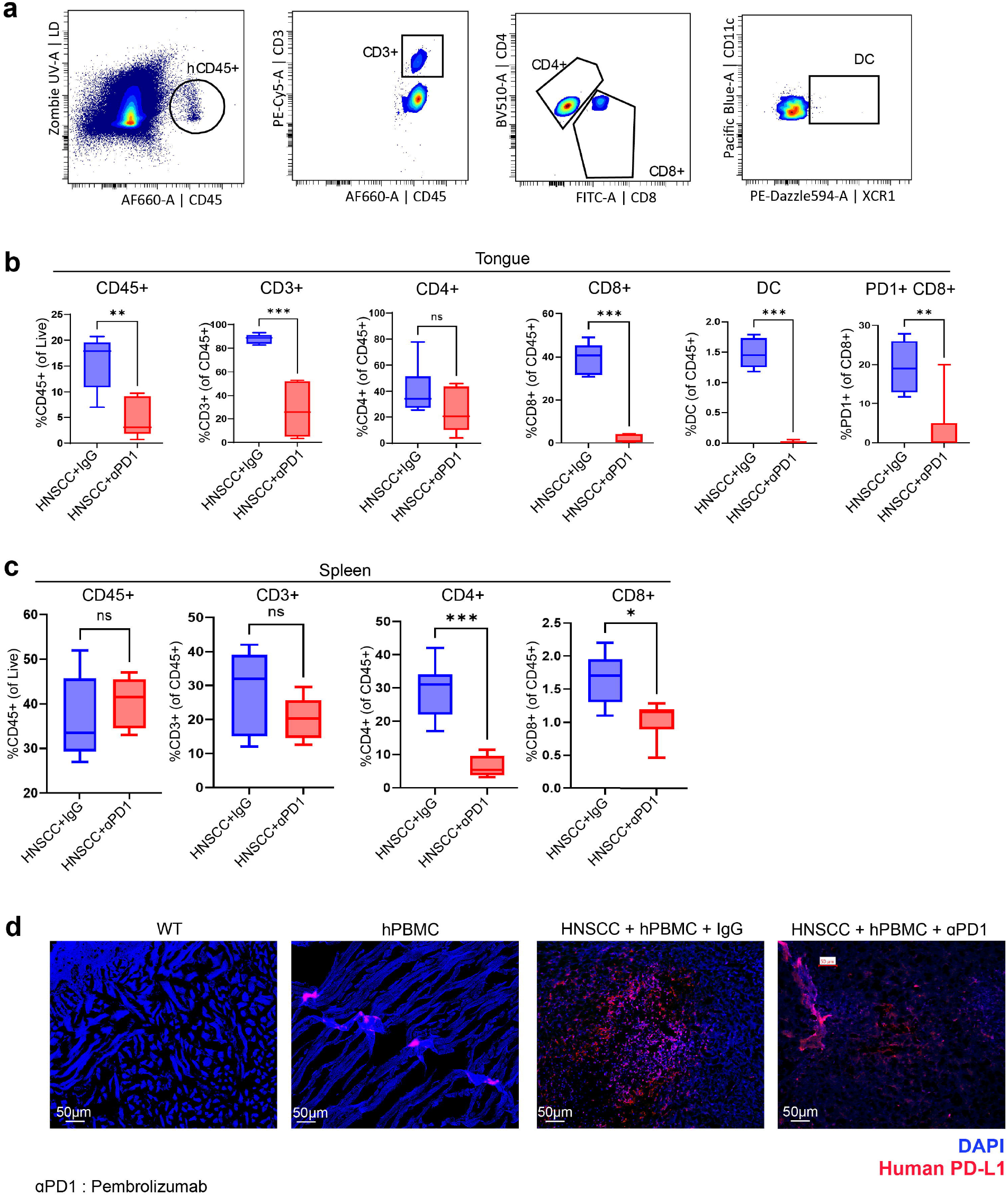
Immune profiling and resistance validation following Pembrolizumab treatment. **a)** Flow cytometry gating strategy showing altered human immune cell populations in the tongue tumor microenvironment after Pembrolizumab treatment. **b)** Quantitative analysis of human immune cell infiltration in the tongue reveals a reduction in key effector populations, including CD8+ T cells and dendritic cells, following treatment. **c)** Assessment of immune cell populations in the spleen shows diminished proliferation and systemic immune suppression post-treatment. **d)** Immunofluorescence staining of tongue tissue demonstrates elevated expression of human PD-L1, indicating an immunosuppressive tumor microenvironment and potential mechanism of therapeutic resistance

Programmed Death Ligand 1(PD-L1) expression in cancer cells indicates an immune evasion mechanism to protect cancer cells from CD8 + T cell-mediated cytotoxicity.^23^ Hence, we evaluated the levels of human PD-L1 in tumor sites using immunofluorescence staining (Fig. 4d). This finding suggests an adaptive immune resistance mechanism at the tumor sites. The upregulation of PD-L1, in conjunction with T cell depletion, indicates a tumor-driven immunosuppressive environment that may blunt the efficacy of checkpoint blockade in αPD-1-treated mice.

### Validation of human transcriptomic landscape in αPD-1 resistant humanized mice

To further characterize the molecular landscape in αPD-1 the resistant humanized OSCC mouse model, we performed bulk RNA sequencing of tongue tumor tissues followed by DGE analysis by aligning the reads to the human genome (hg38) (Table S2). This mapping of the human genome validates the integration of human cells, which enables detailed profiling of therapy-induced variations in gene expression. A heatmap of differentially expressed genes revealed distinct transcriptional signatures between the αPD-1 treated mice and IgG-treated controls (Fig. 5a). Gene set enrichment analysis (GSEA) identified several significantly altered biological processes in the αPD-1 group, including *Positive Regulation of Cell Migration, Extracellular Matrix Degradation*, and *Cell Junction Organization* (Fig. 5b). Canonical pathway analysis further highlighted the key signaling pathways (human-specific) altered in αPD-1 resistant humanized mice (Fig. 5c). Pathways such as *Ephrin A Signaling, PTEN Signaling, Integrin Signaling*, and *Ferroptosis Signaling* exhibited significant activation or inhibition, as indicated by z-scores and -log(*p-*values) (Fig. 5c). Finally, qPCR validation of the human-specific immune markers *HLA-A* and *HLA-B* (Fig. 5d, e) confirmed the transcriptional activity of hPBMCs within the tumor microenvironment, thereby enhancing the translational accuracy of the model. Together, these transcriptomic and pathway enrichment analyses provide mechanistic insights for understanding resistance to anti-PD-1 therapy and highlight the utility of this model for dissecting immune-tumor interactions at the system level.

**Figure 5:**
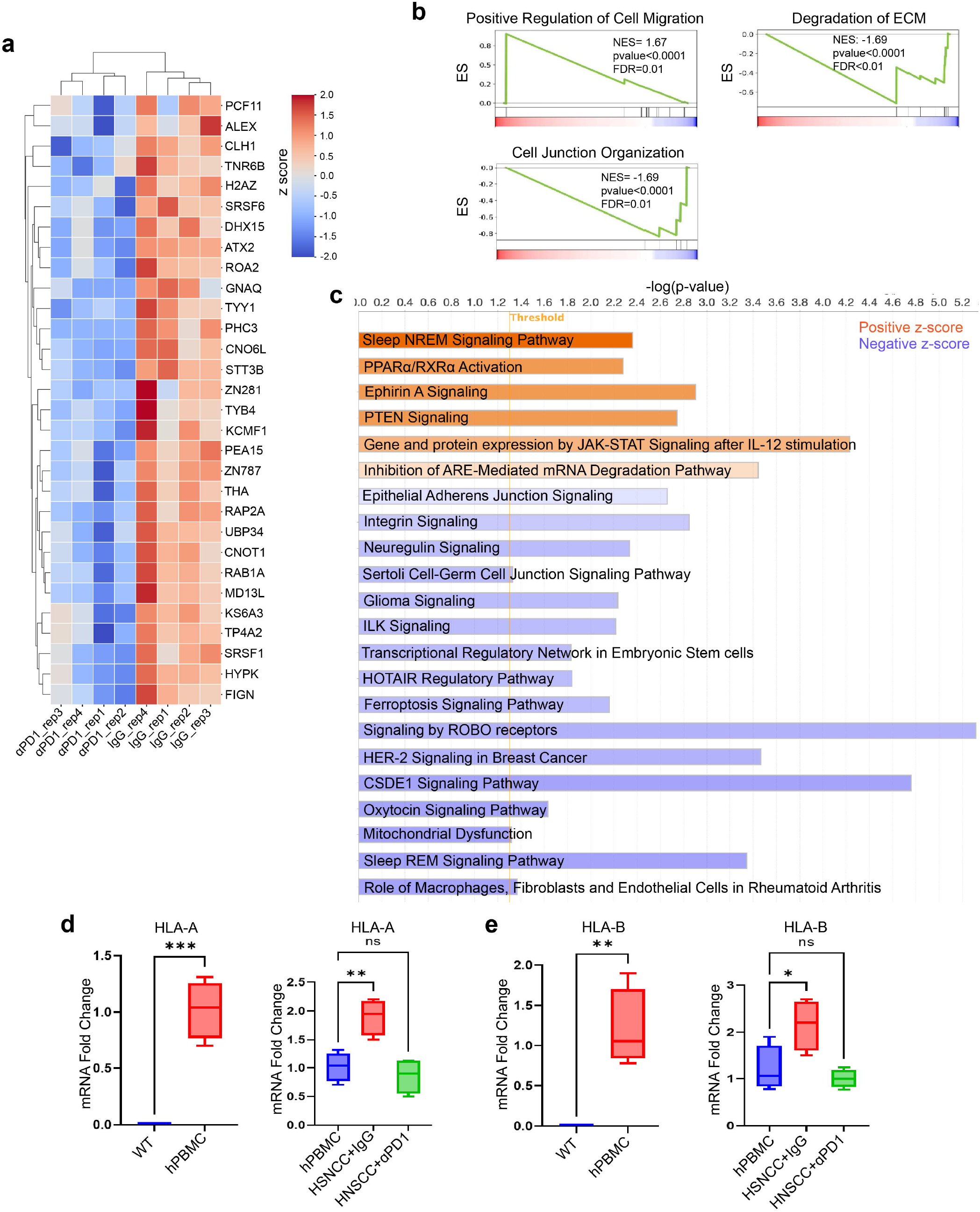
Transcriptomic and pathway alterations validation in humanized mice treated with anti-PD-1 immunotherapy. **a)** Heatmap of differentially expressed genes across αPD1 resistant humanized mice compared to the IgG controls. **b)** GSEA plots show altered gene sets based on normalized enrichment scores (NES). Pathways include *Positive Regulation of Cell Migration, Extracellular Matrix Degradation*, and *Cell Junction Organization*. c) Canonical pathway analysis showing both activated and inhibited pathways (human specific) based on z-scores. Bar length represents z-score magnitude; statistical significance is shown as –log(*p*-value). **d-e)** Quantitative PCR analysis of human HLA-A and HLA-B mRNA expression in PBMCs from wild-type (WT) controls and humanized mice bearing HNSCC tumors, treated with either control IgG or anti-PD-1 antibodies. Bar plots show relative fold changes normalized to WT controls.

## Discussion

This study presents a robust and clinically relevant humanized mouse model to study anti-PD-1 resistance in OSCC. By engrafting hPBMCs into NCG mice and implanting HNSCC stem cells orthotopically, we developed a model that mimics the key aspects of human OSCC, such as aggressive tumor growth, immune cell infiltration, and metastatic dissemination. This dual-engraftment method, which combines human immune cell reconstitution with orthotopic implantation of primary OSCC cells, allows for the investigation of human immune-tumor interactions and treatment responses in a physiologically adequate *in vivo* setting. Unexpectedly, treatment with pembrolizumab led to enhanced tumor progression, increased metastatic burden, and reduced overall survival in humanized mice bearing HNSCC tumors. These paradoxical effects suggested the emergence of immune-mediated resistance mechanisms in this model.

A notable and therapeutically important finding was that pembrolizumab had a paradoxical impact on tumor promotion. Similar to the irAEs observed in patients, treated mice showed increased tumor burden, increased metastasis, and decreased survival.^13,14^ Flow cytometry analysis revealed a decrease in CD8 + T cells, along with a reduction in DC and downregulated MHC-I expression, suggesting a breakdown in antigen presentation and cytotoxic T cell function. These findings align with the clinical data linking CD8 + T cell loss and DC dysfunction to poor immunotherapy outcomes.^24–27^

CD8 ^+^ T cells are crucial for effective antitumor immunity primarily because of their cytotoxic activity. A reduction in CD8^+^ T cells is associated with poor clinical outcomes in cancer patients undergoing anti-PD-1 therapies because of impaired cytotoxic function and diminished tumor control.^24,28^ DCs play a pivotal role in antigen presentation and T-cell priming; hence, their depletion significantly compromises adaptive immune responses, fostering tumor immune evasion.^25,27^ Additionally, MHC-I expression is critical for antigen recognition by CD8^+^ T cells, and its downregulation is frequently observed in resistant tumors, directly contributing to immune escape mechanisms.^26^

Despite its strengths, however, this model has several limitations. However, it does not fully capture the complexity and heterogeneity of human immune responses or the tumor immune microenvironment. Nonetheless, it offers a powerful platform for dissecting resistance mechanisms and evaluating novel therapeutic combinations in a controlled and humanized context.

In summary, our findings highlight the complexity of immune dynamics in response to anti-PD-1 therapy and underscore the utility of this humanized model in uncovering resistance relevant to clinical outcomes. This humanized OSCC model represents a significant methodological advance in preclinical immunotherapy and HNSCC research. This lays a foundation for the mechanistic investigation of resistance pathways and provides a translational bridge to accelerate the development of more effective immunotherapeutic strategies for OSCC and other solid tumors.

## Methods

### Humanized NCG-HNSCC Stem Cell Mice Model

The NCG mouse strain (NOD-Prkdc^em26Cd52Il2rg^em26Cd22/NjuCrl) was generated by sequential CRISPR-Cas9-mediated editing of the *Prkdc* and *Il2rg* loci on an NOD/Nju genetic background. This coisogenic, triple-immunodeficient model lacks functional T, B, and natural killer (NK) cells and exhibits impaired macrophage and dendritic cell functions, making it highly permissive for human immune system engraftment.

Since NCG mice are immunodeficient and carry a mutation in the SIRPα gene, they allow efficient engraftment of foreign hematopoietic cells. Prkdc knockout induces a severe combined immunodeficiency disease-like (SCID-like) phenotype, impairing T-cell and B-cell formation. Il2rg knockout further exacerbated the SCID-like phenotype and reduced NK cell production. These mice can host xenograft cells, tissues, and human immune system components, making them ideal for studies in oncology, immunology, tumor biology, infectious diseases, graft-versus-host disease (GvHD), diabetes, regenerative medicine, hematopoiesis, and tissue transplantation. The NCG mouse model can also be humanized by engrafting peripheral PBMCs or CD34 stem cells to closely mimic the human immune system.

In this study, NCG mice were intravenously injected via the tail vein with 1 × 10L human PBMCs (hPBMCs) per mouse. Humanization was performed using the PBMC Select Humanization Kit (Charles River Laboratories), according to the manufacturer’s protocol for GvHD-compatible PBMC engraftment (https://criver.widen.net/s/mxqcjbqgpx/rm-ps-ncg-pbmc-select-humanizationkit).

### Orthrotopic HNSCC implantation

Three days after engraftment, the mice were anesthetized and orthotopically implanted with 2.5 × 10L patient-derived HNSCC stem cells (Cellprogen) (Patient/donor details are in Supplymentary Table 1) into the anterior tongue using a standardized protocol (n = 10 per condition). Seven days after tumor implantation, the mice were randomized into treatment groups and administered either IgG isotype control or pembrolizumab (10 mg/kg intraperitoneally, once weekly for 3 weeks) at the tumor site. Mice were monitored throughout the treatment period, and euthanized on day 28 post-implantation. Tongue and spleen tissues were harvested for downstream analysis by flow cytometry and immunofluorescence.

### Cell Authentication

Human head and neck cancer stem cells (Cat# 36125-52P, Lot# 2111210; Celprogen) were authenticated prior to use. Species-specific PCR analysis confirmed the human origin of the cell line, thereby confirming the absence of cross-species contamination. Cell line authentication was performed by Celprogen, and the details are provided in Table S1.

### Flow Cytometry

Flow cytometric analysis was performed using a 5-laser 64-color Cytek Aurora Spectral Flow Cytometer (Cytek Biosciences). A comprehensive list of antibodies and fluorochromes is provided in Table S1. Spectral unmixing was performed using SpectroFlow software (Cytek Biosciences).

T cells were defined as CD3+TCRβ+ subsets within the live CD45+ population and further classified into CD4+ (T helper cells) and CD8+ (cytotoxic T cells). Human-specific markers, including CD45 and CD3, were used to distinguish leukocytes and lymphocytes, respectively. DCs were identified using CD11c cell surface markers. The detailed immune cell markers are documented in Table S1.

### RNA Sequencing and Gene Set Enrichment Analysis

Bulk RNA sequencing was performed by the Novogene Corporation to generate paired-end FASTQ files. Quality control of the raw sequencing data was conducted using FastQC, followed by adapter and low-quality base trimming using Trimmomatic^16^. Cleaned reads were aligned to the human reference genome (hg38) using HISAT2^17^ and gene-level read counts were quantified using featureCounts^18^.

Differential gene expression (DGE) analysis was performed using the DESeq2 package^19^ in R, which applies negative binomial generalized linear models to estimate fold changes and statistical significance. Genes with *p*-values < 0.05 were considered significantly differentially expressed.

Heatmaps and hierarchical clustering of gene expression were generated using standard Python libraries, including pandas, seaborns, matplotlib, and scipy. Gene Set Enrichment Analysis (GSEA) was conducted using the GSEApy package^20^, leveraging hallmark and canonical pathway gene sets from the Molecular Signatures Database (MSigDB).^21^

### Ingenuity Pathway Analysis

Ingenuity Pathway Analysis was performed using IPA software (Qiagen).^22^ We have used RNAseq DGE data were generated with DESeq2 (R/Bioconductor software) to perform the analysis. The genes were included with the criteria of p value <0.05, and Z score ≥ ±0.25.

### Pathological Characterization and Immunostaining

Tongue tissues were harvested from mice at the endpoint, fixed in 10% neutral-buffered formalin, embedded in paraffin, and sectioned at 5 μm thickness. Hematoxylin and eosin (H&E) staining was performed on three sections per mouse (n=3 mice per group).

For immunofluorescence analysis, tissue sections were stained with anti-human HLA-A (MHC-I; APC-conjugated) and anti-human PD-L1 (CD274; PE-conjugated) antibodies (Table S1). Staining was performed on the matched sections from each group to assess the expression of immune markers. Fluorescence images were acquired using a confocal microscope under identical exposure settings across the groups. Quantification of marker expression was performed using ImageJ software (NIH), with the signal intensity and positive cell counts normalized to the total number of DAPI-stained nuclei.

### Statistical Analysis

Data analysis, including dimensionality reduction and clustering, was performed using OMIQ software (Dotmatics). Statistical analyses for boxplots were performed using unpaired two-tailed Student’s *t*-tests and one-way Analysis of Variance (ANOVA). DESeq2 applies Wald’s test to calculate the p-values for differentially expressed genes. Graphical representations, including box plots, were generated using GraphPad Prism software.

## Acknowledgment

The authors acknowledge NIH/NIDCR grants R01 DE031413 and CTSA pilot grant UL1TR001430 to Manish V. Bais.

## Ethical Approval

All experiments were performed with prior approval from the Institutional Animal Care and Use Committee (BUMC IACUC protocol # PROTO201800279) of Boston University.

## Competing interests

The authors declare no conflicts of interest or financial interest regarding the content of this manuscript.

## Author contributions

Amit Kumar Chakraborty, Chumki Choudhury, Rajnikant Raut and Manish V. Bais performed the experiments and analyzed the data. Manish V. Bais, Amit Kumar Chakraborty, Chumki Choudhury helped with data interpretation. Amit Kumar Chakraborty, Rajnikant Dilip Raut and Manish Bais did manuscript editing. Manish Bais conceived of and designed the study, interpreted the data, and wrote the manuscript.

## Availability of data and materials

The authors declare that data supporting the findings of this study are available within the paper and its supplementary information files.

## Notes

### Competing Interest Statement

The authors have declared no competing interest.

